# Distinguishing Photoacoustic and Photothermal Neuron Stimulation Through Quantitative Mapping Spatiotemporal Field Evolution

**DOI:** 10.64898/2026.06.02.729635

**Authors:** Deming Li, Andrew Cheng, Xinrui Gong, Haoxin Huang, Yueming Li, Yifan Zhu, Edward A. Nelson, Ji-Xin Cheng, Guo Chen, Chen Yang

**Affiliations:** Department of Electrical and Computer Engineering, Boston University, Boston, MA, USA; Department of Biomedical Engineering, Boston University, Boston, MA, USA; Department of Chemistry, Boston University, Boston, MA, USA

**Keywords:** photoacoustic, photothermal, neural stimulation, non-genetic neuromodulation, nanomaterial

## Abstract

Neuromodulation is a rapidly advancing strategy for modulating brain function and treating neurological disorders, yet the mechanisms of optical neural stimulation remain incompletely understood. A central challenge is that photoacoustic (PA) and photothermal (PT) effects are typically generated simultaneously and evolve on overlapping spatial and temporal scales, making their contributions to neuronal activation difficult to distinguish. Here, we overcome this limitation by integrating two carbon-based emitters with a spatial-offset pump-probe imaging platform to distinguish PA and PT neuromodulation through direct mapping of spatiotemporal field evolution. The two emitters produce nearly identical thermal fields while exhibiting a 26-fold difference in acoustic output, enabling decoupled comparison of thermal and acoustic contributions. To directly characterize the physical fields, we establish a spatial-offset pump-probe imaging system capable of visualizing both temperature and pressure evolution, confirming matched thermal profiles between the two emitters. Neuronal stimulation experiments further show that neurons are markedly more responsive to PA stimulation than to PT stimulation. Beyond resolving a longstanding mechanistic question, this work demonstrates the enhanced efficiency of PA stimulation and provides general guidance for designing high-performance optical neural interfaces.

## Introduction

High-precision neuromodulation serves as an essential tool for understanding the function of neural systems and an alternative potential treatment of drug-resistent neurological disorders ^1–3^. Among non-genetic neuromodulation approaches, light-based methods are particularly promising because they offer safe, versatile and targeted neural control with high spatiotemporal resolution ^4–7^. However, traditional optogenetics faces limitations in penetration depth and raises safety concerns when applied to human ^8^. To overcome these challenges, novel approaches such as Photothermal (PT) and Photoacoustic (PA) neuromodulation have emerged. Photothermal neuromodulation, also referred to as infrared neurostimulation (INS) ^9–11^, activates neurons through localized temperature changes. However, the associated thermal effects limit its applications in clinical treatments, including potential tissue damage and off-target heating ^12,13^. Photoacoustic stimulation provides a distinct alternative, inducing neuron response via the mechanical effect without the thermal toxicity ^14,15^. By applying light-absorbing materials on the interface, it can convert nanosecond laser pulses into rapid thermal expansion and compression, generating ultrasonic waves to modulate neuron activities with single-cell resolution and superior biocompatibility^16–19^.

On the way to understanding the mechanism of PT and PA-induced neurostimulation, the field has systematically explored various mechanistic hypotheses. For PT neuromodulation, both direct thermal elevation and rapid temperature changes serve as effective stimulation mechanisms. Albert et al. reported that infrared heating induced a local temperature rise of ∼10 °C, which evoked TRPV4-dependent voltage responses and action-potential firing in sensory neurons ^11^. In parallel, Lyu et al. have shown that local heating can activate TRPV1-expressing neurons and trigger action potentials ^20^. Apart from a large temperature increase, neurons can also be stimulated through rapid temperature changes. Shapiro et al. demonstrated that a rapid temperature rise can excite neurons by changing membrane capacitance and generating a displacement current, which is proportional to the temperature-changing rate rather than the absolute temperature ^9^. For the mechanism of PA neuromodulation, studies revealed that mechanosensitive ion channels are involved. When mechanosensitive ion channels are overexpressed, the ultrasound stimulation efficiency is boosted ^14,21^. Similar to PT modulation, ultrasound could also stimulate neurons through modulating the membrane capacitance. Shoham group ^22,23^ proposed a biophysical model to calculate how the temperature change and ultrasound waves affect the membrane capacitance. This theory explained both PA and PT neurostimulation mechanisms, that photothermal effects deform the membrane via thermal-induced expansion of the lipid bilayer, whereas photoacoustic effects deform the membrane via mechanical intramembrane cavitation.

Although PA and PT modulation share similar mechanisms, they are always coupled together during PA generation. Previous work demonstrated that only less than 3% of the energy is converted into ultrasound, and the majority of energy turns into heat through a photothermal process ^24,25^. Essentially, PT is inevitable when PA is generated. Therefore, which effect plays the dominant role in neural activition and which one has a lower stimulation threshold, has not yet reached a decisive and exclusive conclusion. Experimental efforts have been made to solve this debate. Hongchae B. et al. ‘s work^26^ demonstrated that a combination of mechanical and thermal effects has a higher success rate than pure mechanical focused ultrasound (FUS) stimulation in evoking mouse tail movements. To separate the two effects, our group previously utilized a nanosecond pulsed laser for PA stimulation and a continuous-wave (CW) laser for PT stimulation to compare the induced neuron activities ^21^. The results indicated that PA stimulation requires ∼30–40 times lower energy than PT stimulation in cultured neurons, suggesting a potentially safer and more efficient stimulation modality. However, since PT stimulation may arise from two different mechanisms, this comparison only ruled out activation driven by the absolute temperature increase. It remains an ongoing debate whether the accompanying thermal effect contributes to the stimulation via a rapid temperature rise mechanism.

To study this essential topic systematically, an existing challenge is that the temperature rise profile produced by a nanosecond pulsed laser evolves on nanosecond timescales of rapid change and micrometer scales of spatial wavelength, making it difficult to capture with conventional probes. Therefore, a method capable of mapping out the relevant physical fields with high spatiotemporal resolution is needed to better understand the mechanism of PA and PT neuromodulation. To this end, a valuable imaging technology published by our group previously, spatial-offset pump-probe imaging (SOPPI) ^27^, offers the capability to simultaneously and quantitatively compare PA and PT field evolution. By spatially offsetting the pump and probe beams, SOPPI detects refractive-index changes induced by both ultrasound and heat. The system offers nanosecond temporal resolution, micrometer spatial resolution, and up to 65 MHz detection bandwidth, while maintaining a millimeter-scale field of view. These capabilities make SOPPI perfectly suited for visualizing both fast acoustic transients and slow thermal diffusions in a broad range of biological systems and serve as an ideal platform for quantitatively distinguishing PA and PT contributions in neuromodulation.

In this work, we innovatively integrate two tools: a pair of PA emitters and the SOPPI imaging system to distinguish PA and PT contributions in neuromodulation. As schematically shown in Fig. 1, an efficient PA emitter produces both a temperature increase and a strong ultrasound field, whereas a weak PA emitter generates an identical temperature field but much weaker acoustic pressure under the same laser excitation. To quantitatively compare not only the total temperature increase and ultrasound pressure, but also the temperature changing rate, we used SOPPI to map the spatial-temporal evolution of PA and PT fields in both emitters, with nanosecond and micrometer resolution. The two emitters were applied in vitro for neurostimulation, and it shows that the efficient PA emitter robustly stimulates neurons, whereas the weak PA emitter elicits a negligible response, indicating that neuronal activation under these conditions is primarily driven by PA rather than PT effects. This unique combination provides a direct experimental strategy to decouple PA and PT effects, which has not been previously achieved to our knowledge. More broadly, the finding not only deepens our mechanistic understanding of PA neuromodulation but also offers design insights for next-generation optical biointerfaces with improved precision, efficiency, and safety.

**Figure 1.**
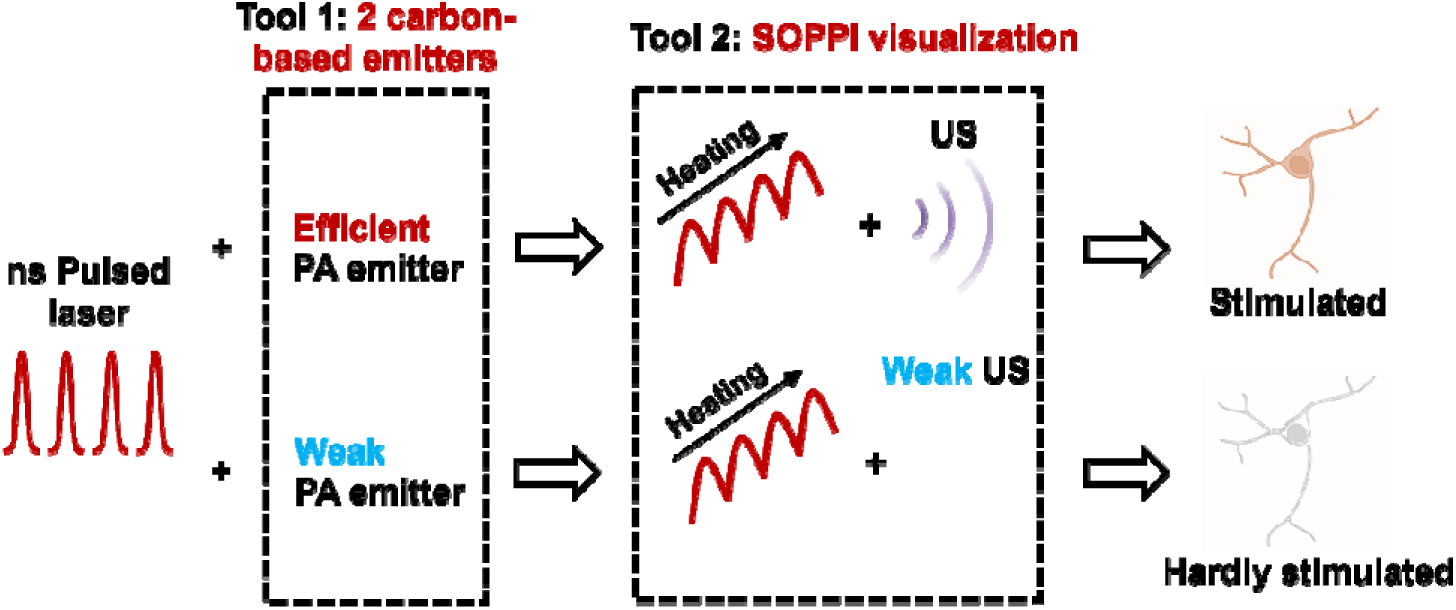
Decoupling PA and PT contributions to neuromodulation using paired carbon-based PA emitters and SOPPI visualization.

## Results

### Fabrication of CSFOE and CNTFOE

In this study, we chose candle soot-based fiber optoacoustic emitters (CSFOE)^28^ and carbon nanotube-based fiber optoacoustic emitters (CNTFOE)^29^ as the platform to study the neuron response to controlled photoacoustic and photothermal stimulation. CSFOE, with an absorber of candle soot, functions as an efficient photoacoustic emitter, whereas CNTFOE is less efficient with an absorber of low concentration carbon nanotube. Importantly, under the same optical excitation, the two emitters can produce comparable photothermal effects but generate markedly different acoustic outputs. This contrast makes the two devices ideal for isolating and comparing photothermal and photoacoustic mechanisms.

To fabricate the efficient PA emitter, candle soot (CS) and polydimethylsiloxane (PDMS) were selected as the key coating materials. Candle soot exhibits high light absorption coefficient and low interfacial thermal resistance, making it an outstanding choice compared to other carbon materials^30^. PDMS possesses high thermal expansion, enabling it to expand and compress efficiently to temperature changes. Combining these two materials creates an exceptionally efficient PA emitter suitable for neuromodulation. The fabrication of CSFOE is shown in Fig. 2a. A 200 μm optical fiber is inserted into a fiber ferrule and placed into the center of a candle flame near the wick for 5 seconds. The interface of the fiber tip and the flame will combust insufficiently and generate flame-synthesized candle soot. The coated candle soot acts as the absorption layer, and its thickness is controlled to 39.28 ± 3.53 μm (N = 8 samples) by the coating time (Fig. S1). PDMS is then deposited on the CS-coated fiber with a thickness of 20-50 μm by a 3D manipulator. Since PDMS penetrates the soot layer, the interface forms a CS-PDMS mixture. After 100°C curing PDMS for 1h, a representative photo of a CSFOE is shown in Fig. 2c.

**Figure 2.**
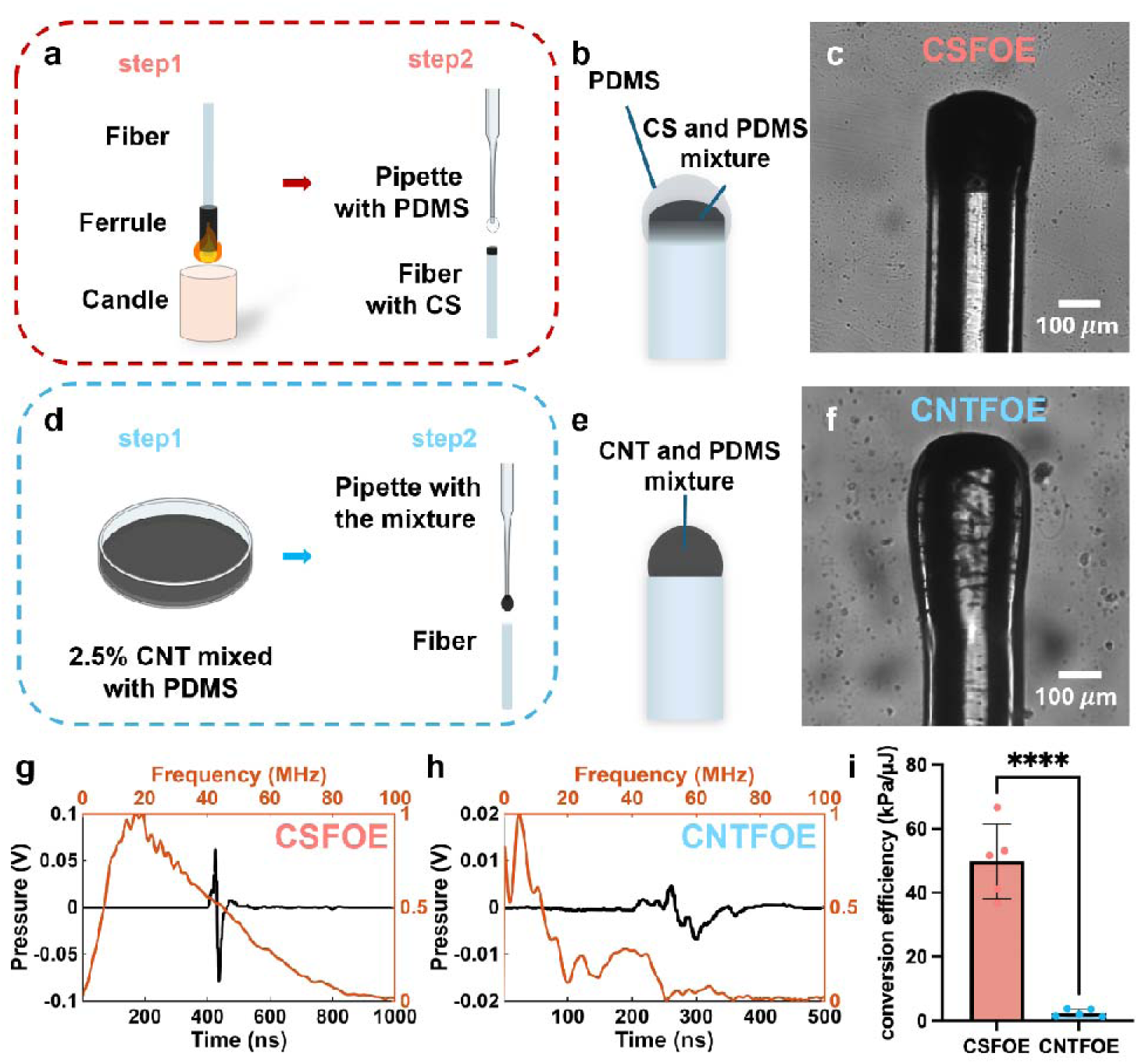
Fabrication and characterization of CSFOE and CNTFOE. (a) Key steps of CSFOE fabrication. (b-c) A schematic and a photo of CSFOE. (d) Key steps of CNTFOE fabrication. (e-f) A schematic and a photo of CNTFOE. (g-h) The generated PA signal and frequency analysis of (g) CSFOE and (h) CNTFOE. (i) PA conversion efficiency (peak-to-peak pressure over laser pulse energy) of CSFOE and CNTFOE (N=5 samples). T-test, **** p < 0.0001.

To fabricate the weak PA emitter, multi-wall carbon nanotube (CNT) was selected as the absorption layer. As demonstrated by Shi et al., the absorber concentration can be modified to tune the generated ultrasound intensity^31^. Here, we choose a relatively low concentration of 2.5%, aiming for a PA emitter with low conversion efficiency while maintaining the consistent photo thermal heating. The fabrication of CNTFOE is shown in Fig. 2d. CNT is mixed with PDMS at a concentration of 2.5% by weight. The tip of a 200 μm multimode optical fiber is carefully cut to get a smooth end. A pipette on a 3D manipulator is used to deposit the mixture onto the tip of the fiber, as Fig. 2d shows. After 100°C curing for 1h, an example of CNTFOE is shown in Fig. 2f.

### PA Characterization of CNTFOE and CSFOE

To compare the PA performance of CNTFOE and CSFOE, a 1030 nm nanosecond pulse laser with a pulse width of 3 ns and a repetition rate of 2.5 kHz is used to generate the photoacoustic wave. The laser energy delivered to the CNTFOE is 12.29 μJ/pulse, while the laser energy delivered to the CSFOE is 4.08 μJ/pulse. The generated ultrasound wave is measured by a hydrophone in water, and the distance between the hydrophone and both FOE tips is ∼ 300 μm (Fig. S2). It can be seen clearly in Fig. 2g-h that the PA signal profile by CSFOE is obviously narrower and stronger than that of CNTFOE. The CSFOE generated a peak-to-peak pressure of 210.34 kPa, while the CNTFOE only generated a peak-to-peak pressure of 15.58 kPa under 3 times higher incident power. The frequency for the fiber optoacoustic emitters was calculated using the fast Fourier transform and is plotted in red in Fig. 2g-h. The CNTFOEs show a central frequency of 6.99 MHz, while CSFOEs show a central frequency of 18.72 MHz, approximately 3 times higher than that of the CNTFOEs. This is because the CNT/PDMS mixture has a relatively lower optical absorption coefficient compared with CS, which would lead to a longer effective absorption depth and therefore result in a broadened frequency, according to the Beer–Lambert law^32,33^. Statistical analysis revealed a significant difference in PA conversion efficiency (p < 0.0001), with CSFOE exhibiting 51.5 kPa/μJ compared to CNTFOE’s 2.4 kPa/μJ. The 21.5-fold difference demonstrates that CSFOE is a much more efficient PA emitter, enabling CSFOE and CNTFOE an ideal platform for direct comparison of PA contributions to neural activation.

### SOPPI Mapped PA and PT fields generated by FOE

Though the PA signals are characterized by hydrophone, quantitative mapping of the PA and PT signals generated by the two FOEs over different distances needs to be done for a systematic understanding and comparison. To directly visualize the distribution of thermal and ultrasound fields generated by the two carbon-based emitters, a custom-built spatial offset pump-probe imaging system (SOPPI)^27^ was established for the measurement (Fig. 3a). A nanosecond pulse laser (Ekspla, NT 252) with a wavelength of 1030 nm, a repetition rate of 20 Hz, a pulse width of 4 ns, and an output power of 46 mW was delivered to the FOE as the pump beam to generate pressure. The tip of the FOE was fixed in a glass plate filled with water. A 1310 nm CW laser (1310LD-4-0-0, Aero DIODE Corporation) was used as a probe laser and focused at the plane of the fiber tip through a 10× water-dipping objective. The generated thermal and ultrasound fields will conduct a refractive index change of water, which can be sensed by the probe laser and detected by the photodiode. A translation stage (ProScan III, Prior) was used to map the generated PA/PT fields with a step size of 8 μm.

**Figure 3.**
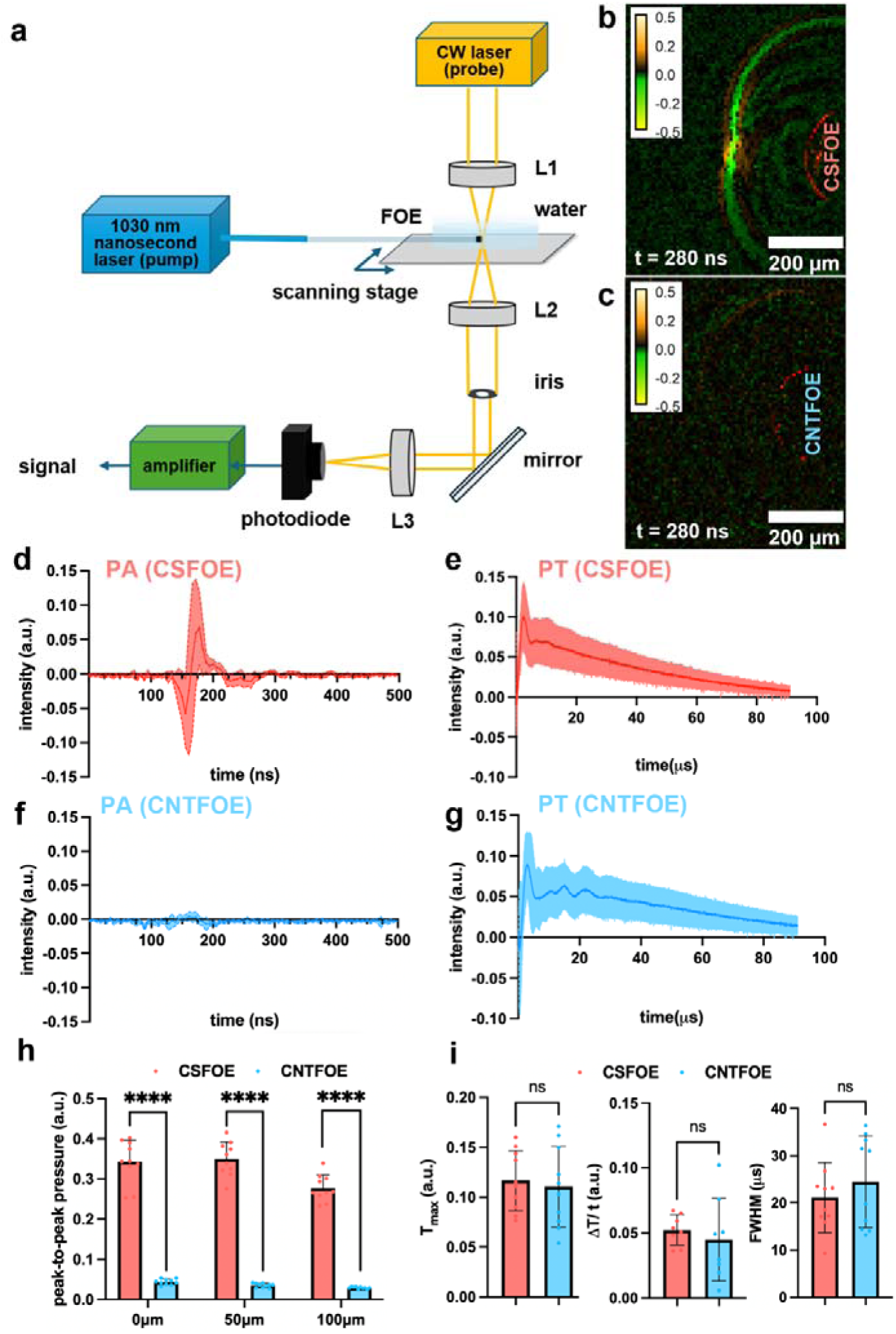
PA and PT field measurements by SOPPI. (a) Schematic of SOPPI system. (b-c) PA (orange and green) and PT fields (red) mapped by SOPPI at *t* = 280 ns. *t* = 76 ns is the pump laser on. Position of FOEs are labelled in the figure correspondingly. (d-g) Averaged PA signal at the distances of 100 µm and PT signal at the FOE surfaces, i.e. distance = 0. (d-e) generated by CSFOE, and (f-g) generated by CNTFOE. N = 10. The shaded areas are standard deviations. (h) Comparison of maximum pressure signals of CSFOE and CNTFOE at the distance of 0 µm, 50 µm and 100 µm. N = 10. T-test, **** *p* < 0.0001. (i) Comparison of thermal signal (total thermal increase, rise rate (total thermal increase divided by the rising time), and the temporal width of thermal field (FWHM)) of CSFOE and CNTFOE. T-test, **** *p* < 0.0001, ns, no significant difference, *p* = 0.7160, 0.5502, 0.3962, respectively.

Representative signals mapped by SOPPI at the tip of CSFOE and CNTFOE at t = 280 ns were shown in Fig. 3b and Fig. 3c, respectively. The thermal field was located at the tip of FOEs, while the pressure propagated forward as a spherical wave. The average of 10 PA traces measured 100 µm away from the CSFOE can CNTFOE tips were plotted, and the generated PA pressure showed a bipolar signal. As shown in Fig. 3d and Fig. 3f, CSFOE generated a 16 times stronger pressure than CNTFOE under the same laser energy. Notably, here time *t* = 76 ns is the laser on time, which means the PA waves would start to propagate after that. To further compare the propagation of PA waves, the peak-to-peak pressure at the distance of 0 µm, 50 µm and 100 µm were compared in Fig. 2h. Within the 100 µm propagation, the pressure experienced an average attenuation of 19.28%. All the t-tests between the peak-to-peak pressure generated by CSFOE and CNTFOE showed a significant difference (*p* < 0.0001), which is consistent with the pressure measured by the hydrophone.

At the same time, the thermal traces measured at the tip of FOEs were averaged and plotted in Fig. 2e-g as well. Both CSFOE and CNTFOE showed a rapid increase within 3 µs followed by slow decay. Statistical analysis of total thermal increase (T_max_), thermal rise rate (ΔT/Δt), and temporal width of thermal pulse (FWHM) were conducted in Fig. 3i. No significant differences were observed for CSFOE and CNTFOE in peak thermal intensity (0.114 vs. 0.108 a.u., p = 0.7160), thermal rise rate (0.046 vs. 0.043 a.u./µs, p = 0.5502), or temporal width of the thermal field (23.43 vs. 24.17 µs, p = 0.3962). These results demonstrated that CSFOE and CNTFOE generate identical PT fields along with significantly different PA fields, which means that they can serve as a comprehensive experimental framework for this decouple PA and PT contributions to neuron stimulation.

### More effective neuron stimulations induced by CSFOE than CNTFOE

The SOPPI measurements offer us clear evidence that CSFOE and CNTFOE will generate 16 times different pressure under the same laser condition, while maintaining the identical thermal fields. This result guides us to use the two FOEs to decouple PA and PT contributions in neuromodulation. GCaMP6f-labeled primary neurons (DIV 10-13) were cultured on a glass-bottom dish, and calcium imaging was performed to monitor neuronal activity. The position of FOEs was precisely controlled by a 4D micromanipulator to approach the target neurons and control the distance from the fiber tip to the neurons (Fig. S3). A 3 ns pulsed laser train at 1030 nm with a repetition rate of 2.5 kHz, a duration of 100 ms, a laser pulse energy of 16.4 μJ/pulse, and an energy density of 39.9 mJ/cm^2^ was delivered to the FOE. The distance between the FOE tip and the neuron is controlled as close as possible without touching, and regarded as 0 μm. It can be clearly observed that CSFOE was able to induce the neuron activity (Fig. 4a), while for the CNTFOE, no calcium response can be seen at the same laser condition and distance (Fig. 4b). The calcium responses of neurons to CSFOE or CNTFOE stimulation were plotted as a function of time in Fig. 4c-d (N=20 for CSFOE and N=17 for CNTFOE). For CSFOE, all the cell traces show clear responses after the laser is on, to a greater or less extent. A few of them were even over stimulated, and didn’t return to the baseline within 10 secs. However, CNTFOE induced negligible calcium responses for all the neurons.

**Figure 4.**
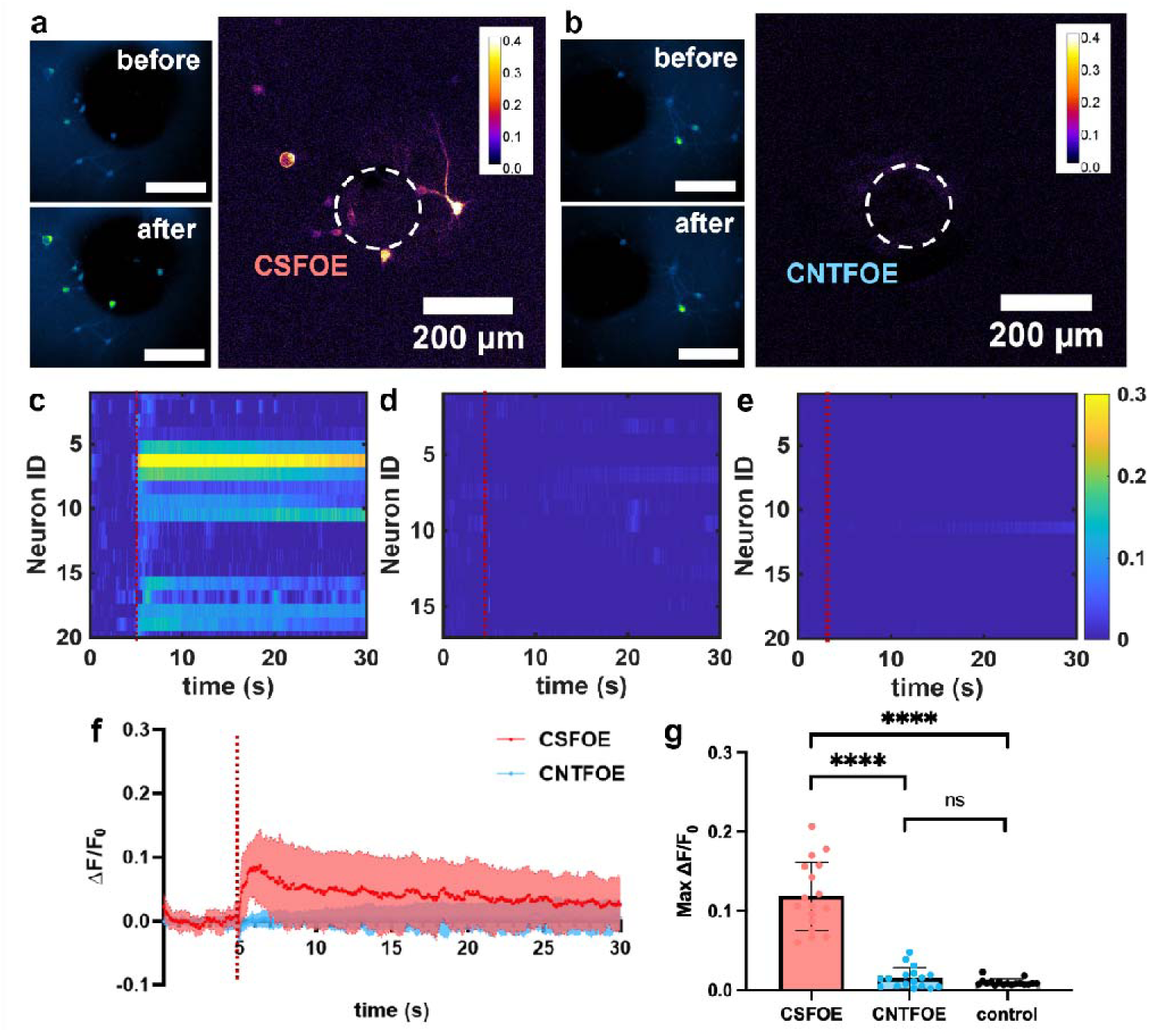
Neuron activation induced by CSFOE and CNTFOE. (a-b) Image of the maximum fluorescence change induced by (a) CSFOE and (b) CNTFOE stimulation. Laser energy: 41 mW, tip-to-neuron distance: 0 µm, duration: 100 ms. (c-e) Colormap of (c) CSFOE stimulation, N = 20; (d) CNTFOE stimulation, N = 17, (e) laser only control, N = 20. Red dashed line: laser on time, *t* = 5s. (f) Average calcium traces of neurons of CSFOE and CNTFOE stimulation. Red dashed line: laser on time. (g) Comparison of the maximum fluorescence change of CSFOE stimulation, CNTFOE stimulation, and laser only control. t-test, **** *p* < 0.0001, ns, no significant difference, *p* = 0.1177.

A laser-only control experiment was done under the same laser condition with a bare optical fiber, and no activation was observed, as shown in Fig. 4e. The result demonstrated that the modulation of neural activities was from the photoacoustic coating, but not from the laser directly. Moreover, the averaged traces of fluorescence change of the two FOEs were plotted in Fig. 4f. The trace of CSFOE exhibited a rapid increase upon laser onset and returned to the baseline afterwards, which indicated that this is a safe condition for neurons. The t-test results showed that the average maximum ΔF/FL of 11.87% in the CSFOE group is significantly higher than in both the CNTFOE and control groups (*p* < 0.0001 for both comparisons). In contrast, no significant difference is observed between the CNTFOE and control groups (*p* = 0.1177).

These experiments demonstrated that two FOEs has two drastically different neuromodulation responses, and CSFOE stimulates neurons more effectively than CNTFOE under the same laser condition and same distance towards neurons. During this process, there are three different mechanisms related to this non-genetic modulation as mentioned in the introduction: (1) neuron being stimulated through mechanical force; (2) through temperature sensitive ion channels triggered by the total temperature increase; (3) the membrane capacitance change triggered by the rapid change of temperature. By combining SOPPI measurements, we can not only compare the total temperature change, but also confirm the temperature changing rate of the two FOEs are identical, which comprehensively demonstrated that the two FOEs generated identical thermal fields. Since the identical PT fields of CSFOE and CNTFOE would contribute equally to the neural activation, the reason (2) and (3) didn’t dominate to the efficient neuromodulation induced by CSFOE. Thus, we can conclude that it is the significant difference in PA that dominate these different stimulation results. This finding filled the gap in our previous study^21^ that only compared the contribution of absolute temperature change with PA, and completely dispelled the doubts about whether PA actually played a role in neuron stimulation.

### Matched neuron stimulation results with PA distribution

To thoroughly compare the PA-induced and PT-induced neuromodulation, a laser condition that is strong enough for CNTFOE to stimulate neurons is needed. Here, the laser power was increased to 51.8 mW, and the duration was increased to 200 ms. The distance between the tips of FOEs and the neuron was controlled to be 0 _μ_m, 30 _μ_m, 50 _μ_m, 100 _μ_m, and 150 _μ_m to obtain different intensities of PA and PT fields. As is shown in Fig. 5a, CSFOE can overstimulate the nearby neurons under this strong laser condition. When the CSFOE is moved to 50 μm, although the fluorescence change remained the same, the calcium traces had a gradual down-forward trend, indicating this is a relatively safer condition. When the CSFOE further moved to 100 μm, the calcium fluorescence change was obviously weaker than the previous distances, which would be relevant to the PA attenuation during propagation. For CNTFOE under the same laser condition, it can induce neuron responses at distances of 0 μm and 50 μm as well. However, the induced fluorescence change at 100 μm is almost negligible. To better compare the stimulation results, the maximum fluorescence change at different distances was measured and plotted. It can be seen from Fig. 5g that 0 μm, 30 μm, and 50 μm shared a similar fluorescence change, the average of which is 0.201, 0.216, and 0.205, respectively. With the distance increasing to 100 μm and 150 μm, the maximum fluorescence change experienced an obvious decrease, both of which are less than 10%. When it comes to CNTFOE, the average max ΔF/F_0_ only reached 0.07 even at the distance of 0 μm, which is 3 times less than that of CSFOE. Notably, what CSFOE and CNTFOE have in common is the drop beyond 50 μm. The averaged maximum ΔF/F_0_ are 0.041 and 0.036 at the distance of 100 μm and 150 μm, respectively. Therefore, CSFOE can stimulate neurons more effectively than CNTFOE, evidenced by the larger averaged max ΔF/F_0_ at all distances.

**Figure 5.**
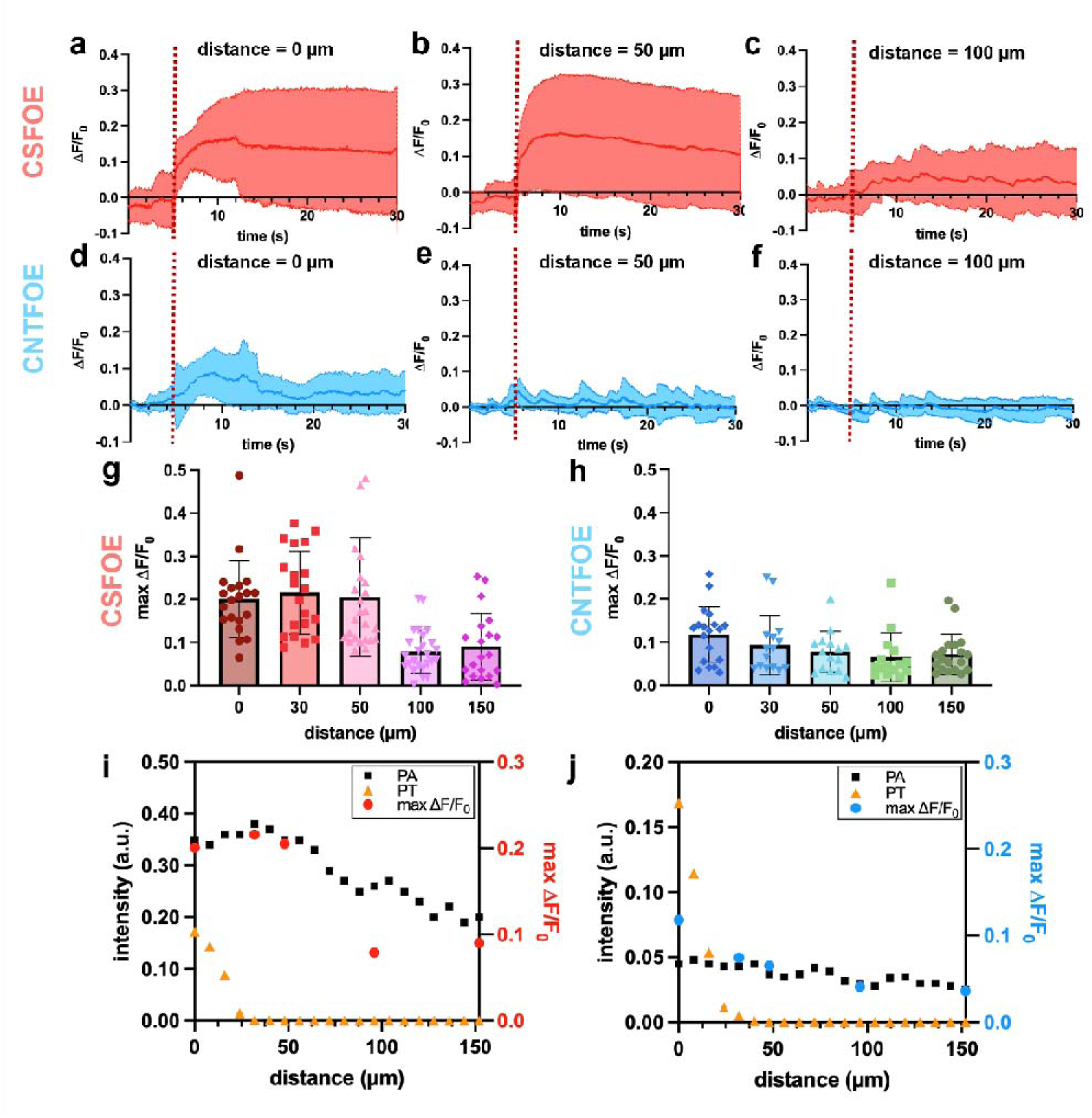
Neuron stimulation induced by CSFOE and CNTFOE at different distances. (a-c) Average calcium traces of neurons of CSFOE stimulation with the tip-to-neuron distance of (a) 0, N = 25, (b) 50, N = 25, (c) 100, N = 20, duration: 200 ms, laser power: 51.8 mW, red dashed line: laser on time. (d-f) Average calcium traces of neurons of CNTFOE stimulation with the tip-to-neuron distance of (d) 0, N = 25, (e) 50, N = 15, (f) 100, N = 20, duration: 200 ms. Red dashed line: laser on time. (g-h) Maximum fluorescence change at different distances induced by (g) CSFOE and (h) CNTFOE stimulation. (i-j) SOPPI measured PA signal, PT signal, and maximum fluorescence change obtained from Calcium imaging as a function of distance under (i) CSFOE and (j) CNTFOE stimulation.

Critically, to further confirm our hypothesis that it is PA that contribute to the neuron stimulation, the peak-to-peak PA intensity, maximum PT intensity measured by SOPPI, and average maximum fluorescence change were plotted along the distances in Fig. 5i and Fig. 5j. The pressure of both CSFOE and CNTFOE slightly increased around the distance of 30 μm, resulting from the focusing effect of the PDMS layer^27^. After that, due to the water absorption, the PA intensity gradually decreased with the distance. The PT fields localized at the FOE tip, and dropped rapidly within 20 μm. As we can see clearly, the average max ΔF/F_0_ of CSFOE perfectly suits the PA intensity. It is the PA attenuation that is responsible for the weaker neuromodulation beyond the distance of 50 μm. Notably, for CNTFOE, the averaged max ΔF/F_0_ also matched well with the PA intensity distribution. Within the distance of 30 μm, the combination of PT and PA generated by the CNTFOE activated neuron. While beyond the distance of 30 μm, it may only the weak PA instead of PT that contributed more to the induced neuron activities, since the PT intensity beyond 20 μm was negligible. This systematic study of distance with PA and PT distribution shows the spatial correlation between the PA intensity and stimulation efficiency for the two FOEs. Therefore, we can conclude that PA would stimulate neurons more effectively under these laser conditions.

## Discussion and Conclusion

Photoacoustic (PA) modulation is an emerging technology that can safely activate neurons with high spatial and temporal precision. In previous studies, both in vitro, such as wild-type neurons^28,34,35^, and in vivo, such as in the rodent brain^19,36^ have been shown to respond to these PA signals with single-cell and even subcellular precision. However, since

PT is inevitably coupled with PA during generation, it remains unclear whether PA or PT dominates the neural activation. To resolve this, two carbon-based fiber emitters: candle soot-based fiber optoacoustic emitters (CSFOE) and carbon nanotube-based fiber optoacoustic emitters (CNTFOE) are chosen for this study, which absorb identical heat but generate different pressure under the same laser condition. The PA and PT field generated by the two FOEs was visualized by a custom-built spatial-offset pump-probe imaging system (SOPPI) to compare in detail. By stimulating neurons with equal PT but different PA fields, we could decouple the individual contributions of each mechanism. Results demonstrated that neurons exhibit significantly greater sensitivity to PA than to PT, indicating that PA, rather than the accompanying thermal effect, is the primary driver of neural activation.

The key innovation is that we establish the two kinds of FOE, which generate a greatly different pressure with the same heat absorption. For the perspective of CSFOE, it has such an outstanding performance on producing PA, owing to not only the intrinsic material advantage of candle soot, but also its optimized double-layered design. Prior work showed that candle soot/PDMS composites can achieve a markedly 6 times higher photoacoustic efficiency^30^ than CNT composites because the CS/PDMS interface has lower interfacial thermal resistance. This allows heat to transfer more rapidly into PDMS, whose large thermal expansion then generates stronger acoustic waves. In addition, candle soot particles have a three-dimensional ball-shaped nanostructure, while carbon nanotubes have a one-dimensional geometry. Therefore, CS is more favorable for thermal energy release, further reducing thermal loss and enhancing optoacoustic conversion. For the perspective of CNTFOE, the concentration of CNT is controlled to 2.5% instead of the commonly used 10%, to further decrease the pressure conversion efficiency of CNTFOE. Unlike candle soot coating methods, which offer limited control over absorption efficiency (Fig. S1), the CNT concentration in PDMS can be systematically adjusted to precisely tune PA emission. This is feasible because the concentration directly affects the effective absorption thickness of the absorption layer, and further determines the ultrasound generation intensity^31^. As a result, CSFOE achieved a 21.5 times stronger acoustic output than CNTFOE under the same laser incident energy.

To better understand the mechanism of PA- and PT-induced neurostimulation, many efforts have been put in recent years. PA can stimulate cells mainly through a mechanical effect. When a short laser pulse is absorbed by the photoacoustic material, the resulting rapid thermal expansion generates a localized acoustic pressure wave. This pressure wave deforms the nearby cell membrane, which can activate mechanosensitive ion channels, alter membrane capacitance, and perturb membrane-associated proteins^14,15,37,38^. For PT neural stimulation, two main mechanisms have been proposed. The first is a direct thermal effect, where light-induced temperature increases of several degrees activate thermosensitive ion channels^39,40^. The second is an opto-capacitive mechanism, in which a rapid temperature rise modulates membrane capacitance and induces transmembrane capacitive currents^9,41,42^. Guo Chen et al.‘s work^21^ used a nanosecond laser and a CW laser to compare PA-induced neuromodulation with the direct thermal effect, which is the first mechanism. Here, our work bridged the gap of comparing PA with the accompanied PT mechanism, including the maximum thermal rise, thermal rising rate and the temporal width of thermal field. The results showed that it is PA that contributes more to the neuron responses in this process.

However, several limitations should also be considered. Firstly, the experiments were performed under a specific in vitro platform, and the results may not fully capture the complexity of in vivo neural tissues, where scattering and network-level interactions could alter the stimulation outcome. Secondly, because of the limited laser pulse energy available, it remains unclear how many times of optical energy would be required for the CNTFOE to evoke the same fluorescence change as CSFOE, and whether that ratio would match the PA conversion efficiency ratio between them. A more rigorous quantitative comparison will require future experiments with an expanded laser energy range. Besides, since the present comparison is between two stimulation platforms with matched PT fields and various PA fields, a purely PT condition without any accompanying PA component was not directly established. Thirdly, the two FOEs produced acoustic signals with different central frequencies. As is shown in Fig. 2g-h, CSFOE generates the pressure with a ∼3 times higher frequency than that of CNTFOE, and it remains unknown whether neurons will respond differently to different frequencies and therefore represent a potential confounding factor beyond pressure amplitude alone.

In conclusion, this work provides direct experimental clues on distinguishing the relative contributions of PA and PT effects in neurostimulation. More broadly, this work helps clarify the physical mechanism of non-genetic neuromodulation and offers a useful framework for comparing different stimulation strategies. By distinguishing the roles of PA and PT mechanisms, this study also provides guidance for safe, efficient, and highly localized tools for both basic neuroscience research and therapeutic applications.

## METHODS

### FOE fabrication

Both CSFOE and CNTFOE were made from a 200 μm multimode optical fiber (FT200EMT, Thorlabs, Inc., NJ, USA). Before the coating process, the fiber tip was carefully cut to get a smooth surface, and the cross-section was checked via an edge microscope.

For CSFOE, the tip of the optical fiber was inserted into a fiber ferrule and flushed with it. Then the ferrule with fiber was placed into the center of a paraffin wax candle flame near the wick for 5 seconds, until it became fully coated with candle soot. Polydimethylsiloxane (PDMS) was prepared by gently dispensing the silicone elastomer (Sylgard 184, Dow Corning, USA) into a container to reduce air entrapment, followed by mixing with the curing agent at a 10:1 weight ratio. The prepared PDMS was subsequently applied to the CS coated fiber tip using a nanoinjector. During this process, both the fiber and the nanoinjector were positioned with a 3D micromanipulator to ensure accurate alignment, and the coating procedure was monitored in real time using a laboratory-built microscope. The coated fiber was then cured in an incubator at 100°C for one hour.

For CNTFOE, multi-wall CNTs (<8Lnm OD, 2−5Lnm ID, Length 0.5−2 μm, VWR, Inc., NY, USA) was mixed with PDMS prepared following the above procedure, at a concentration of 2.5 % by weight. The mixture was carefully dripped onto the fiber tip using a 3D micromanipulator, and the alignment was monitored in real time using a lab-made microscope. The coated fiber was then cured in an incubator at 100°C for one hour.

### Characterization of photoacoustic properties of the FOEs

An acoustic characterization of the generated optoacoustic waves was conducted by a lab-made hydrophone system. A compact Q-switched diode-pumped solid-state laser (1030 nm, 3 ns, 100 μJ, repetition rate up to 10 kHz, RPMC, Fallon, MO, USA) served as the excitation source. The laser was first connected to an optical fiber and then connected to the FOE with a fiber optic attenuator set (multimode, varied gap of 2/4/8/14/26/50 mm, SMA Connector, Thorlabs, Inc., NJ, USA) to adjust the laser power. For acoustic pressure measurements, a 85-µm needle hydrophone (HGL-0085, Onda Corporation, USA) was employed together with an ultrasonic pre-amplifier (0.2–40 MHz, 40 dB gain, Model 5678, Olympus,USA). The distance from the FOE tip to the hydrophone was maintained at 300 μm using a 3D micromanipulator. Signals from the hydrophone were collected using a digital oscilloscope (DSO6014A, Agilent Technologies, CA, USA). All components were immersed in deionized water to emulate conditions relevant to biomedical applications and to minimize acoustic impedance mismatch among the FOE, water, and detectors. The frequency data was obtained through the Fast Fourier Trans-form (FFT) using MATLAB 2020a.

### Mapping the PA and PA field by SOPPI

A spatial-offset pump-probe imaging system (SOPPI) was applied to map the PA and PT field evolution. A 1030 nm nanosecond pulse laser (Ekspla, NT 252) with a repetition rate of 20 Hz, a pulse width of 4 ns, and an output power of 46 mW was delivered to the FOE as the pump beam. A continuous wave 1310 nm laser (1310LD-4-0-0, AeroDIODE Corporation) serves as the probe beam, with a power of 5 mW, and was used to detect the signal. The probe laser was sent to a water dipping objective (UMPLFLN 10XW, Olympus) for illumination and collected by an air objective (MPlanFLN 10×, Olympus). The signal-carrying probe laser was detected by an amplified InGaAs photodiode (PDA05CF2, Thorlabs) with a 1310 nm bandpass filter (FBH1310-12, Thorlabs). The output signal was connected to a 50-ohm resistor, amplified by a 46-dB amplifier (100-MHz bandwidth, SA230F5, NF Corporation), and digitized by a data acquisition card at 180 MSa/s (ATS9462, Alazar Tech). A translation stage (ProScan III, Prior) was used to scan the fiber that delivered the pump laser and the generated PA/PT fields with a step size of 8 μm. The data collection was triggered by both the pump pulse and the translation stage. The time t = 0 ns is the trigger signal sent time, and it takes ∼76 ns for the laser to receive the signal and be turned on. Each pixel corresponds to a single pump pulse.

### Neuron culture

All experimental procedures complied with all relevant guidelines and ethical regulations for animal testing and research established and approved by Institutional Animal Care and Use Committee (IACUC) of Boston University (PROTO201800534). Primary cortical neurons were isolated from embryonic day 15 (E15) Sprague−Dawley rat embryos of either sex (Charles River Laboratories, MA, USA). Cortices were isolated and digested in TrypLE Express (ThermoFisher Scientific, USA). Then the neurons were plated on poly-D-lysine (50 µg mL−1, ThermoFisher Scientific, USA)-coated glass bottom dish (P35G-1.5-14-C, MatTek Corporation, USA). Neurons were first cultured with a seeding medium composed of 90% Dulbecco’s modified Eagle medium (ThermoFisher Scientific, USA) and 10% fetal bovine serum (ThermoFisher Scientific, USA) and 1% GlutaMAX (ThermoFisher Scientific, USA), which was then replaced 24 h later by a culture medium composed of Neurobasal Media (ThermoFisher Scientific, USA) supplemented with 2% B27 (ThermoFisher Scientific, USA), 1% N2 (ThermoFisher Scientific, USA), and 1% GlutaMAX (ThermoFisher Scientific, USA). On day three, the pAAV.Syn.GCaMP6f.WPRE.SV40 virus (Addgene, MA, USA) was added to the cultures at a final concentration of 1 μL/mL for GCaMP6f expression and has cre-independent expression already with no Cre virus involved in the process. Half of the medium was replaced with fresh culture medium every 3–4 days. Cells cultured in vitro for 10–13 days were used for FOE stimulation experiment.

### In vitro neurostimulation

In vitro neurostimulation experiments were performed using a Q-switched 1,030-nm nanosecond laser (1030 nm, 3 ns, 100 μJ, repetition rate up to 10 kHz, RPMC, Fallon, MO, USA). A 3-D micromanipulator (Thorlabs, Inc., NJ, USA) was used to position the FOE in the cell culture dish. The distance between the tips of FOE and neuron was controlled to be 0 μm, 30 μm, 50 μm, 100 μm, and 150 μm. To label the distance, the tip of FOE is continuously moving down until starting to touch one neuron, and the distance is marked as 0 μm and applied to other neurons. The actual distance ranges from 0 -10 μm; for simplicity, here are labeled as 0 μm.

Calcium fluorescence imaging was performed on a lab-built wide-field fluorescence microscope based on an Olympus IX71 microscope frame with a 10 × air objective (UPLAN FLN 10×, 0.3 NA, Olympus, MA, USA), illuminated by a 470 nm LED (M470L2, Thorlabs, Inc., NJ, USA) and a dichroic mirror (DMLP505R, Thorlabs, Inc., NJ, USA). Image sequences were acquired with a scientific CMOS camera (Zyla 5.5, Andor) at 20 frames per second. Neurons expressing GCaMP6f at DIV (day in vitro) 10–13 were used for the stimulation experiment.

### Data analysis

Prism 10 and MATLAB were used to analyze optoacoustic signal data from the hydrophones. Calcium imaging for Fig. 4 and Fig. 5 were processed using ImageJ, and other calcium imaging analysis including fluorescence intensity traces, statistical analysis, PA and PT intensities were completed with Prism 10.

## Author contributions

GC and CY conceived the idea; DL, AC, and HH fabricated and characterized FOEs; DL and GC designed and performed SOPPI measurements; DL, XG, and HH performed neuron experiments; DL, GC, XG, AC, YL, YZ, NE, CY and JXC discussed and analyzed the data; XG, DL, HH, NE prepared neuron cultures; DL wrote the original draft of the manuscript; GC, YL, XG, YZ, CY, and JXC reviewed and edited the manuscript. CY and JXC supervised the project.

## Acknowledgements

This work is supported by NIH R21NS145121 to CY. The cortical neurons were provided by Hengye Man lab at Boston University.

## Data Availability Statement

The data that support the findings of this study are available from the corresponding author upon reasonable request.

