## Supplemental figure 1-3 for "Distinguishing Photoacoustic and Photothermal Neuron Stimulation Through Quantitative Mapping Spatiotemporal Field Evolution"

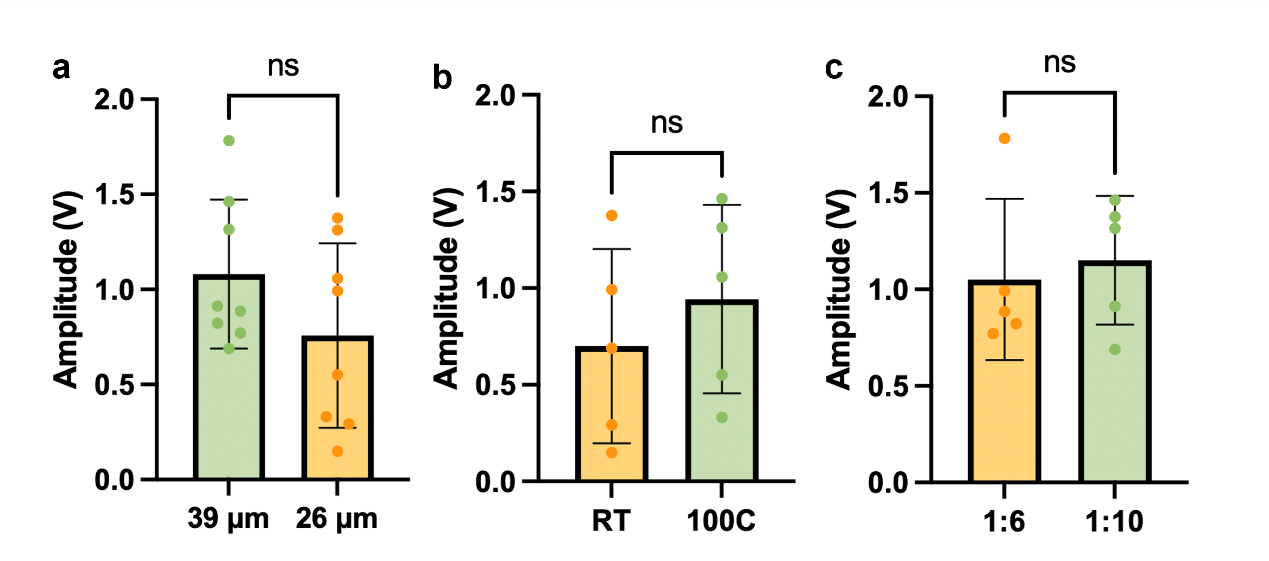

**Figure S1. Peak-to-peak pressure of CSFOE at different conditions.** (a) Thickness of candle soot layer controlled by coating time: 5s: 39 µm $\pm$ 9%; 3s: 26 µm $\pm$18%. T-test, p = 0.1654. N = 8. Standard deviation: 39 µm group: 0.3921; 26 µm group, 0.4848. (b) Curing temperature of room temperature (RT) and 100$℃$(100C). T-test, p = 0.4602. N = 5. Standard deviation: RT group: 0.4872; 100C group: 0.5024 (c) PDMS mixed ratio (curing agent and silicone elastomer base) of 1:6 and 1:10. T-test, p = 6865. N = 5. Standard deviation: 1:6 group: 0.4174; 1:10 group: 0.3333.

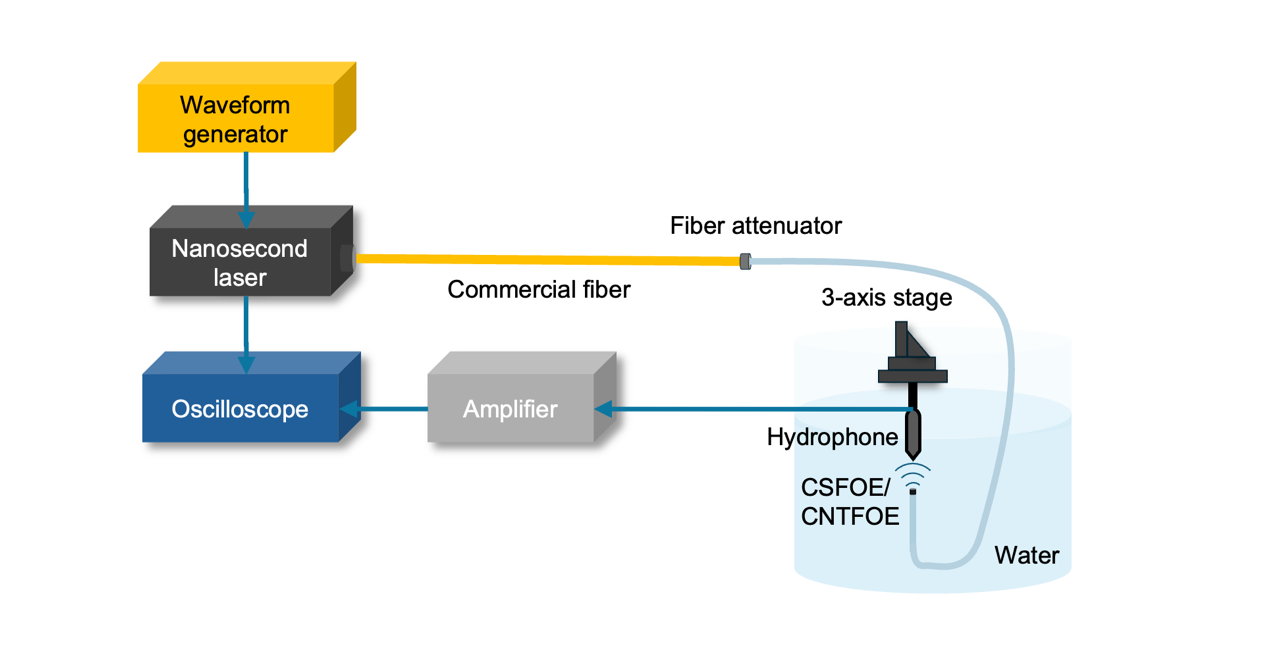

**Figure S2. Schematic of a typical PA measurement.** Waveform generator: square wave; amplitude: 5V peak-to-peak; repetition rate: 10kHz; duty cycle: 30%. Laser: compact Q-switched diode-pumped solid-state laser (RPMC Lasers Inc., USA), wavelength: 1030 nm; pulse width: 3 ns; full output power: 100 μJ; repetition rate: 2.6 kHz. Fiber: multimode optical fiber (Thorlabs, Inc., USA), core diameter: 200 µm; NA: 0.39. Fiber attenuator: varied gap SMA Connector (Thorlabs, Inc., USA). Oscilloscope: DSO6014A (Agilent Technologies, CA). Amplifier: wideband voltage amplifier DHPVA-101 (Femto, DE), gain: 40 dB; bandwidth: DC to 100 MHz.

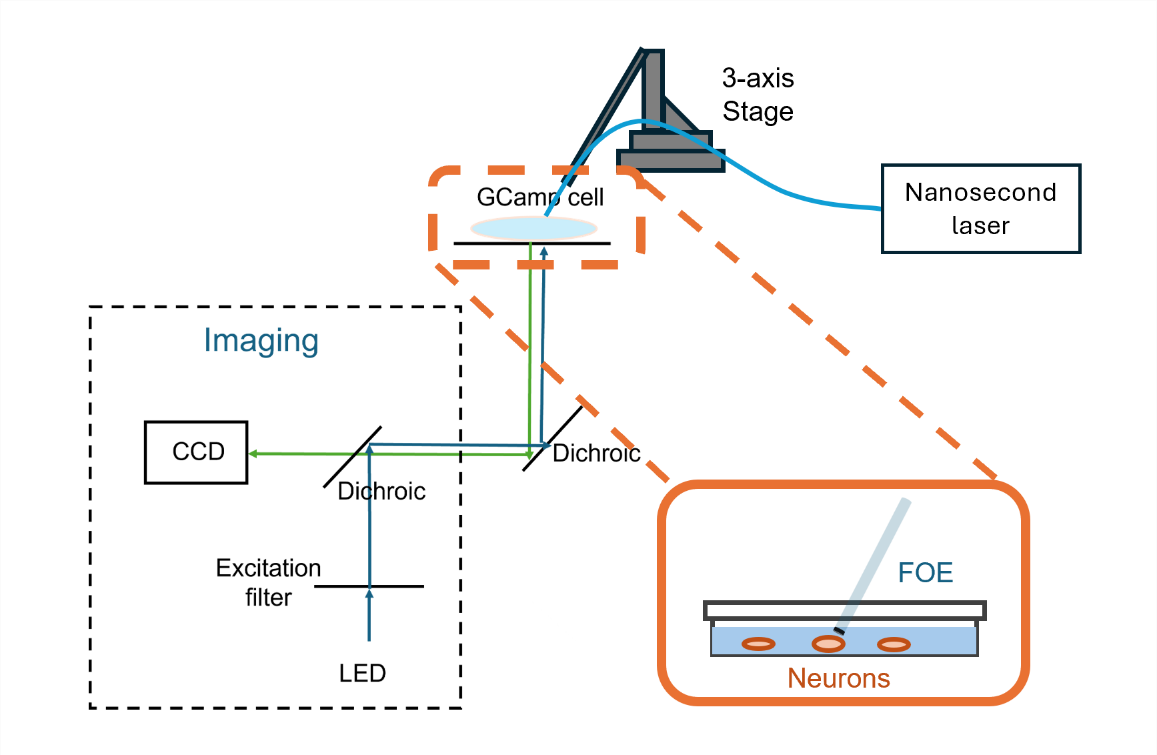

**Figure S3. Schematic of FOE induced neuron stimulation.** In vitro neurostimulation experiments were performed using a Q-switched 1,030-nm nanosecond laser (1030 nm, 3 ns, 100 μJ, repetition rate up to 10 kHz, RPMC, Fallon, MO, USA). A 3-D micromanipulator (Thorlabs, Inc., NJ, USA) was used to position the FOE in the cell culture dish. Calcium fluorescence imaging was performed on a lab-built wide-field fluorescence microscope based on an Olympus IX71 microscope frame with a 10 × air objective (UPLAN FLN 10×, 0.3 NA, Olympus, MA, USA), illuminated by a 470 nm LED (M470L2, Thorlabs, Inc., NJ, USA) and a dichroic mirror (DMLP505R, Thorlabs, Inc., NJ, USA). Image sequences were acquired with a scientific CMOS camera (Zyla 5.5, Andor) at 20 frames per second. Neurons expressing GCaMP6f at DIV (day in vitro) 10–13 were used for the stimulation experiment.
